# Rapid discovery of drug target engagement by isothermal shift assay

**DOI:** 10.1101/584656

**Authors:** Kristofor J. Webb, Kerri A. Ball, Stephen J. Coleman, Jeremy Jacobsen, Michael H.B. Stowell, William M. Old

## Abstract

Identifying protein targets directly bound by drug molecules within living systems remains challenging. Here we present the isothermal shift assay, iTSA, for rapid identification of drug targets. Compared with thermal proteome profiling, a prevailing method for target engagement, iTSA offers a simplified workflow, 4-fold higher throughput, and multiplexed experimental designs with higher replication. We demonstrate application of iTSA to identify targets for several kinase inhibitors in lysates and living cells.

---

A key challenge in drug discovery is the identification of proteins directly bound by small molecule drugs, i.e. target engagement. Many drugs in clinical use are known to promiscuously bind multiple targets in a manner necessary for therapeutic effects, while binding to “off-target” molecules often contributes to toxic side-effects^1–4^. Yet, many drugs approved for human use have not been evaluated for target binding on a proteome scale. For example, an analysis of 243 kinase inhibitors revealed the presence of widespread off-target binding to non-kinase proteins^5^.

Prevailing strategies to determine target engagement exploit the thermodynamic stabilization that occurs due to ligand binding^6^. A notable example is the thermal shift assay^7^, which we have used to optimize membrane receptor stability for subsequent crystallographic studies^8^. Thermal shift assays have been adapted to complex lysates or cells (CETSA)^9^ and allow for proteome-wide assessment of target engagement in a method called thermal proteome profiling (TPP), or MS-CETSA^10^. TPP involves a 10-temperature melting curve and at least two replicates per condition, to estimate the thermal melting temperature (*T*_m_) shift between drug and vehicle for each protein quantified (ΔX in **Fig. 1a**), requiring at least 40 samples and consuming 1-2 weeks of instrument time^10^. This low throughput limits the number of drugs and conditions that can be interrogated, particularly for laboratories without access to dedicated instrumentation.

**Figure 1.**
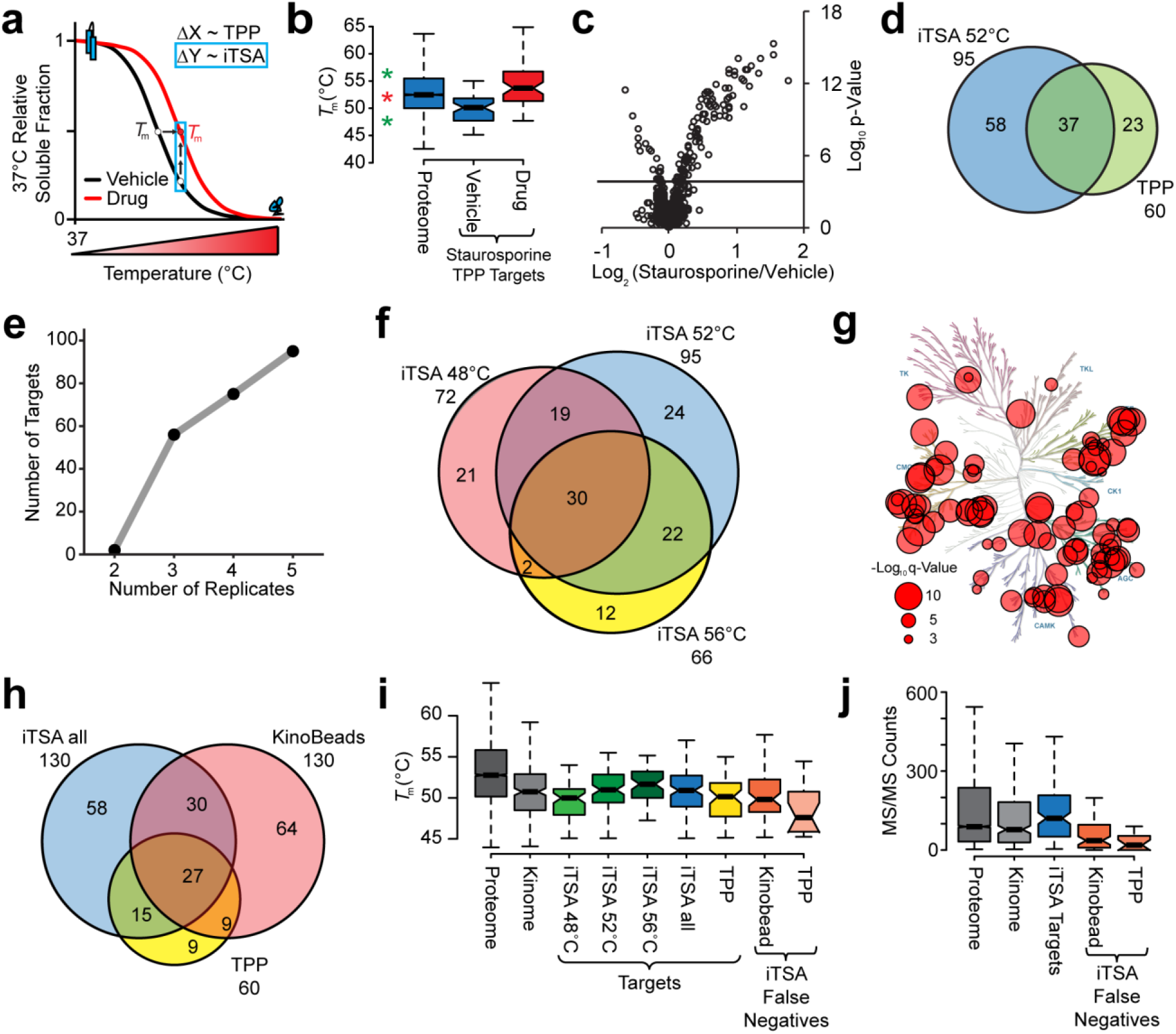
Concept and application of the isothermal shift assay (iTSA) method. **a,** Representative vehicle and drug thermal gradient curves indicating the shift in apparent *T*_m_ upon drug binding, as measured by TPP (ΔX), and the detection of a shift in the melting curves by measurement of differential protein solubility by iTSA (ΔY). **b,** Distribution of Savitski et al.^10^ K562 proteome *T*_m_ (N=5985) and target data (N=60) is shown; the first and third quartiles are spanned by the box and the median shown as the line within the box, notches extend to +/-1.58 IQR/sqrt(n) and whiskers indicating the last data points within 1.5× the interquartile range. **c,** Volcano plot visualization of iTSA performed at 52°C using 20 μM staurosporine. q-value= 0.001 is indicated with solid line. **d,** Venn diagram of iTSA 52°C and Savitski et al.^10^ staurosporine targets. **e,** Number of changes identified with increasing replication, estimated by significance (0.001 FDR) in greater than 80% of sub-sampled combinations from iTSA 52°C. **f,** Targets from 48, 52, and 56°C iTSA experiments. **g,** Kinome tree displaying all iTSA staurosporine kinase targets. Circle size is proportional to the −log_10_ q-value maximum from 48, 52, or 56°C. Illustration reproduced courtesy of Cell Signaling Technology, Inc. **h,** Venn diagram of iTSA targets (from all three iTSA experiments) overlap with previous staurosporine target assays. **i,j,** The distribution of melting temperature (*T*_m_) and MS/MS count was evaluated for the iTSA targets (FDR < 0.001) and compared to the groups indicated using a box-and-whisker plot defined as described in **b** (N in **Supplementary Data File 1**: *boxplot.stats*).

To address these limitations, we reasoned that shifts in thermal stability could be detected by quantifying the difference in soluble protein fraction at a single temperature, carefully selected to maximize detection sensitivity (ΔY in **Fig. 1a**). The resulting isothermal shift assay (iTSA) method requires 4-fold fewer samples and achieves comparable sensitivity to TPP experiments. iTSA simplifies the experimental design and eliminates the need to estimate *T*_m_ shifts. By employing neutron-encoded isobaric labeling, a direct comparison of drug and vehicle replicates can be made within a single multiplexed sample, accommodating up to 5 replicates per condition for increased statistical power.

We selected 52°C for our initial iTSA analysis with staurosporine. Staurosporine, a broad spectrum kinase inhibitor, was an ideal choice for evaluating the proposed iTSA strategy as it allowed benchmarking against targets identified by TPP^10^. As shown in **Fig. 1b** (red star indicating 52°C), this temperature represented the proteome median *T*_m_ and fell in between the vehicle and drug *T*_m_ distributions reported for the TPP identified targets of staurosporine^10^. K562 lysates were treated with vehicle or 20 μM staurosporine then processed using the iTSA workflow as described in **Methods** and illustrated in **Supplementary Fig.1**. We identified 95 proteins with staurosporine-induced shifts in solubility at 52°C (FDR < 0.001), compared with 60 staurosporine targets identified previously by TPP analysis^10^ (**Fig. 1c-d, Supplementary Data File 1**).

The increase in iTSA target identifications over TPP was surprising (95 compared with 60), given that single temperature analysis might be expected to miss proteins with large deviations from the median Tm. Additional analysis of our iTSA data suggested that the increased number of targets may result from the increase in replication (*i.e*. 2 replicates in TPP versus 5 replicates in iTSA). Using the staurosporine iTSA data from our 52°C experiment, we performed sub-sampling to test the effect of replication on the number of identified targets at FDR < 0.001 (**Fig. 1e, Methods, Supplementary Data File 1**). Unsurprisingly, we found that replication improved the sensitivity of the experimental design, and that a single additional replicate (2 vs 3) increased the number of identified targets from 2 to more than 50. The 5 replicate design afforded by TMT 10-plex reagents increased the number of identified targets by nearly a factor of 2, indicating that increased replication beyond that typically used by TPP increases statistical power to detect true hits.

To evaluate the effect of iTSA target temperature selection, we performed iTSA at two additional temperatures: 48°C and 56°C. As shown in **Fig. 1b**, (green stars) these temperatures flanked the interquartile *T*_m_ range of proteome. Surprisingly, both temperatures performed well (**Supplementary Fig. 2a-b**), identifying 72 (48°C iTSA) and 66 (56°C iTSA) staurosporine targets with an FDR < 0.001. iTSA performed at 52°C identified more targets overall and more unique targets. Combining the three temperatures (48, 52, and 56°C) led to a moderate increase in the total number of observed targets, from 95 to 130 (**Fig. 1f**). As would be expected for a kinase inhibitor, the staurosporine targets identified by the iTSA assay were enriched in kinases (**Fig. 1g and Supplementary Fig. 3**).

Next, we performed an orthogonal validation of the iTSA results by comparing the iTSA-identified staurosporine targets to those identified using a kinobead competition binding assay^10,12^. We observed a 47% overlap of iTSA targets with previously identified staurosporine targets, and identified many unique targets (**Fig. 1h**). As expected with a kinase inhibitor, the iTSA targets overlapping with hits from previous studies were predominately kinases. However, we also identified the porphyrin-binding proteins ferrochelatase (FECH) and heme-binding protein 1 (HEBP1), which are not obvious targets of an ATP-competitive kinase inhibitor, but are in agreement with previously reported TPP staurosporine targets^10^. Moreover, the association of FECH with staurosporine and numerous other kinase inhibitors was recently validated.^5,10,15^

Evaluation of the 58 targets uniquely identified by iTSA revealed many more kinases along with kinase-binding proteins and phosphoproteins known to be regulated by staurosporine-targeted kinases (**Supplementary Data File 1**). Kinase-binding proteins would be expected to be indirectly stabilized by staurosporine binding to their cognate binding partner, a phenomenon documented previously^13^. For example, DCAF7 is a stable interaction partner of the kinase DYRK1A^14^; both were identified by iTSA to be strongly stabilized by staurosporine. Likewise, destabilized cognate binding partners have also been noted previously^10^. Consistently, we found that the regulatory subunits of PKA were destabilized by staurosporine, while the catalytic subunits of PKA were stabilized in iTSA.

We also found six additional targets unique to the iTSA method which are not known to bind to staurosporine (RCN2, MIEF2, ALDH6A1, MRPL55, CCDC144C, and Q6ZSR9). Although we can’t rule out that these are false positives, the stringent FDR cutoff of 0.001 predicts no more than one false positive in our set of 130 putative targets. Thus, these represent potential novel staurosporine targets for future validation studies.

One potential caveat associated with the iTSA method is that it failed to identify ~53% of the previously reported targets of staurosporine, with many of these originating from the kinobead study (**Fig. 1h**). Of the kinobead hits, 64 were not identified by iTSA or TPP. To help understand the origin of these false-negatives, we cross-referenced the distributions for *T*_m_ (**Fig. 1i**), and depth of coverage (tandem mass spectra MS/MS counts)(**Fig. 1j**). The *T*_m_ distributions for the targets identified in iTSA experiments (48, 52, 56°C) were surprisingly broad, with medians that increased with the experimental target temperature. The minimum *T*_m_ of staurosporine targets identified by the TPP workflow was lower than for iTSA targets, providing a potential explanation for the few targets that were identified by TPP (TPP False Negatives, **Fig. 1i**). Both TPP and iTSA, however, demonstrated incomplete coverage of the *T*_m_ distribution for known kinases.

All of the workflows demonstrated coverage that extended above and below the interquartile range of both the kinase family and the complete proteome. Moreover, the interquartile *T*_m_ range of the kinobead-derived false negatives was well within the assay range of iTSA, suggesting that the iTSA *T*_m_ range alone could not account for missing these targets. A stronger association with target specificity was observed for depth of coverage (MS/MS count), where the iTSA-identified targets exhibited higher MS/MS counts relative to the false negatives from both orthogonal methods (**Fig. 1j**). This result suggests that these targets were present in lower concentrations relative to proteins identified by iTSA, and thus subject to MS/MS sampling bias inherent in data-dependent acquisition methods. MS/MS sampling could be improved by increased fractionation, technical replication, and increased sequencing speed.

We next asked how iTSA performs using K562 lysates treated with more selective kinase inhibitors, harmine and H-89. Harmine is in an ATP-competitive inhibitor of DYRK family kinases, inhibiting DYRK1A with an apparent Ki of 65 nM^16^. iTSA profiling of K562 lysates with 20 μM harmine revealed stabilization of DYRK1A and its known interacting protein, DCAF7^14^ (**Fig. 2 a,d, Supplementary Data File 2**). Surprisingly, we found that the protein kinases CDK8, CDK9 and CSNK1G3 were stabilized in the presence of harmine. To our knowledge, harmine binding affinities have not been measured for these kinases, but KinomeScan in vitro activity screens found that 10 μM harmine^17^ inhibited CDK8, CDK9 and CSNK1G3 by 79%, 43% and 34%, respectively, indicating that iTSA can detect kinases inhibited more modestly by harmine.

**Figure 2.**
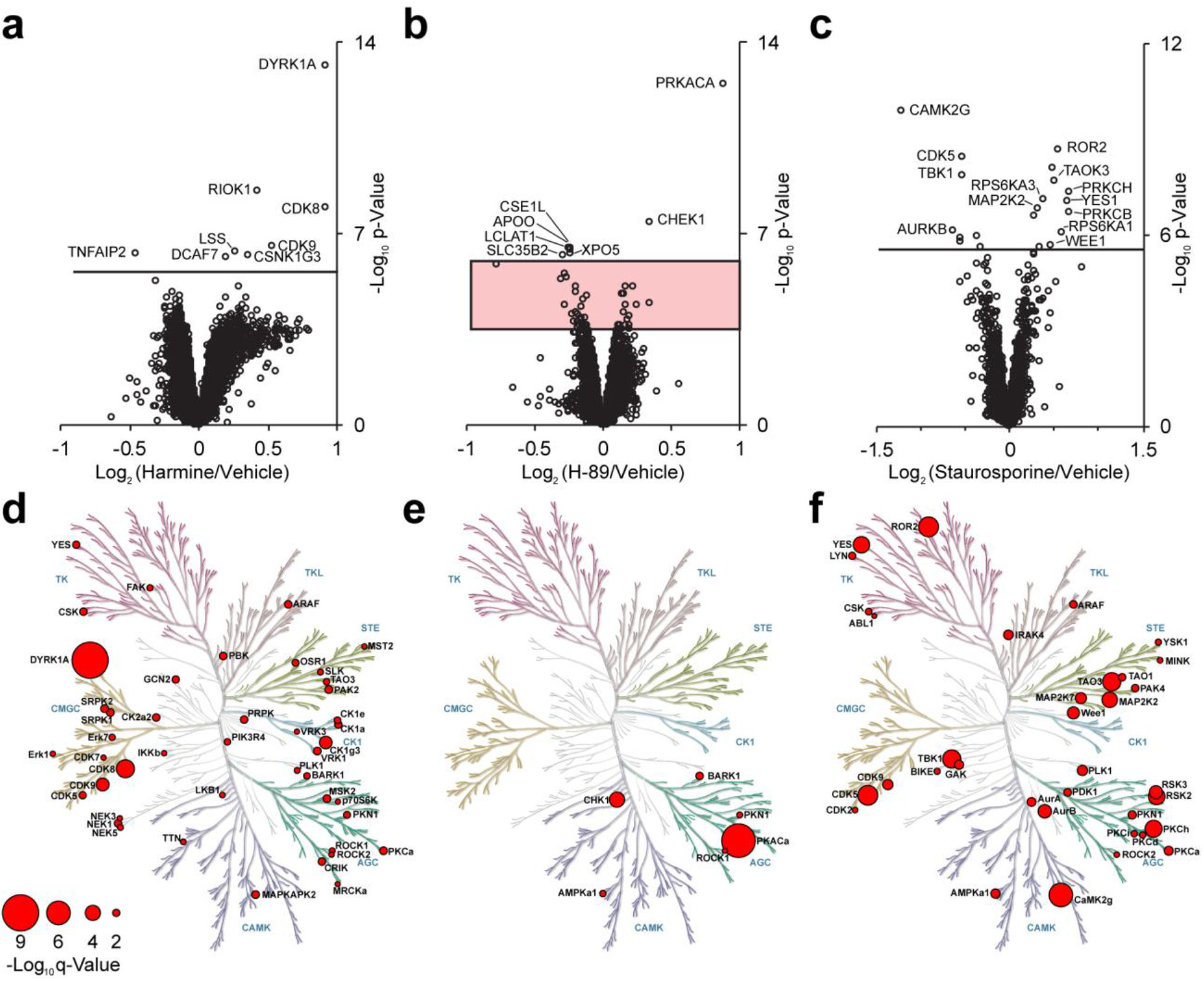
iTSA identifies known kinase inhibitor targets in cell lysates and living cells. Volcano plot visualization of proteins showing changes in thermostability in the presence of three different kinase inhibitors, tested using a two-sample moderated empirical Bayes t-test. FDR = 0.001 is solid line. Red box defines 0.001 > FDR < 0.05. Kinome maps (illustration reproduced courtesy of Cell Signaling Technology, Inc.) display −log10 q-value (size key displayed for all) of kinase targets **a,d,** Using lysate, iTSA was performed at 52°C using 20 μM Harmine. **b,e,** Using lysate, iTSA was performed at 52°C using 100 nM H-89. **b,f,** Using living cells treated with 1 μM staurosporine for 15 minutes, in-cell iTSA was performed at 52°C.

We next examined H-89, an ATP-competitive inhibitor of protein kinase A (PKA). iTSA analysis using 100 nM H-89 in K562 lysates identified the catalytic subunit of PKA (PRKACA) as the most significant target (**Fig. 2 b,e and Supplementary Data File 2**), further demonstrating that iTSA effectively identifies true-positive targets with high confidence. However, here we sought to demonstrate that by lowering the confidence filter (FDR < 0.05 = 95% confidence) iTSA can also be utilized to characterize off-target drug interactions. Using this lower threshold, 107 targets were identified of which only a small percentage were kinases (**Fig. 2e**). This was a significant finding given than the most meticulous characterizations of H-89 off-target binding proteins utilized only purified kinases^18^. Most targets were novel, including several proteins regulating protein import into the nucleus (TGFB1, NUP188, CSE1L, TNPO1, IPO7, TNPO3). While further validation of these novel H-89 off-target binding partners is needed, some of the iTSA low threshold targets have previously been validated, for example, ROCK1^18^. We also identified ATP2A2, which has been reported to bind H-89 in a noncompetitive manner with respect to ATP^19^ and further supports the identification of many non-kinase H-89 binding partners.

To validate iTSA for identification of intracellular target engagement, we performed 52°C iTSA in K562 cells treated with 1 μM staurosporine (**Fig. 2c,f, Supplementary Data File 2**). The lower dose used for intracellular target identification (20 μM for in-lysate iTSA) was necessary to minimize the cytotoxic effects associated with higher staurosporine concentrations^20^. As expected with a lower dose and intracellular ATP concentrations of 1-5 mM^21^, fewer targets were identified (20). Nevertheless, many of the targets identified are kinases (**Fig. 2f**). Only eight of the in-cell iTSA targets, however, overlapped with the in-lysate iTSA, and some targets that were stabilized in the in-lysate iTSA were destabilized in the in-cell iTSA (such as CDK5). The divergence between in-lysate and in-cell target engagement has been documented for MS-CETSA^10^, and is consistent with more complex interactions associated with the primary targets in a cellular environment where ATP and other metabolites are present at physiological concentrations necessary to drive signal transduction.

Here, we have demonstrated that our iTSA workflow provides an attractive method for streamlined identification of high-confidence target engagement in lysates or living cells. The single temperature strategy affords increased sensitivity and throughput relative to traditional approaches. High throughput drug screening is a multi-billion-dollar market^22^. Performed early in the drug development process, iTSA could rule out problematic compounds with undesirable off-target profiles before substantial development investments have been made. Furthermore, iTSA could be used to provide insight into the mechanism of action for compounds identified with phenotypic-screening. iTSA does not require chemical modification and immobilization necessary for molecular-affinity enrichment methods, requires fewer samples and instrument time relative to TPP, and can be accommodated into higher-throughput workflows for identifying target engagement.

## Methods

Methods are available in the Supplementary materials / online version of the paper.

## Acknowledgements

This work was supported by a DARPA cooperative agreement, 13-34-RTA-FP-007, to WMO and MHBS. We thank Dr. Brady Worrell and Dr. Mike Klymkowsky for their technical review and recommendations.

## Author Contributions

MHBS, KJW, KAB, and WMO conceptualized the study. KJW, KAB, SJC & WMO designed the experiments. KJW, KAB, SJC, JJ & WMO designed data analysis strategies. KJW performed the experiments. KAB, JJ, SJC implemented analysis of the data. KAB, KJW, SJC, JJ, MHBS & WMO participated in the interpretation of the data and the writing of the manuscript. WMO and MHBS led the funding acquisition.

## Methods

### Cell Culture

K562 cells were acquired from ATCC. Cells were cultured in RPMI GlutaMAX media containing 10% fetal bovine serum and 1% Anti-Anti antibiotic/antimycotic (Thermo Fisher Scientific, Waltham, MA). Cells were expanded to 1.5 x 10^6^ cells/mL. 50 million cells were washed two times with PBS (137 mM NaCl, 2.7 mM KCl, 10 mM NaHPO4, and 1.8 mM KH_2_PO_4_, pH 7.4) and snap frozen in liquid nitrogen for use in lysate iTSA experiments or were used immediately for in-cell iTSA experiments.

### Preparation of lysate

Frozen cell pellets were suspended in 1 mL of Lysis Buffer (137 mM NaCl, 2.7 mM KCl, 10 mM NaHPO_4_, and 1.8 mM KH_2_PO_4_, 0.4% Igepal CA-630, 1X cOmplete™, Mini, EDTA-free Protease Inhibitor Cocktail (Sigma #11836170001), 1X Pierce™ Phosphatase Inhibitor Mini (Fisher # A32957), pH 7.4). The pellet was suspended by vortex then sonicated for 30 seconds, 3 times, with a 30-second rest in between in a 4°C Bioruptor water bath sonicator (Diagenode, Lorne, Australia). Lysate was centrifuged at 21,000 *g*, 4 °C, for 15 minutes and the supernatant collected. The final lysate protein concentration was then adjusted to 5 mg/mL as determined by BCA assay (Thermo Fisher Scientific, Waltham, MA). Samples were promptly used in thermal profiling experiments.

### Thermal treatment of lysate

To reduce variability, minimal sample handling was a focus of the procedure. For thermal gradient profiling three gradient programs were created using a PTC-200 thermal cycler to cover the temperature points 37, 41.2, 44, 46.8, 50, 53.2, 56.1, 59.1 63.2, 66.9 °C (MJ Research, Reno, NV). Program 1 consisted of temperatures 37, 41.2, 56.1 °C, program 2: 44, 46.8, 50, 53.2 °C, and program 3: 59.1, 63.2, 66.9 °C. The heated lid was set to 95 °C. Three PCR plates were prepared with 40 μL per well of lysate and sealed (individually sealable and removable wells; 4titude Random Access, PN 4ti-0960/RA 96-well plate). The plates were spun at 500 *g* for two minutes at 4 °C, and then kept at 4 °C prior to use (less than 15 minutes). For each program, the thermal cycler program was set to run for a total of 20 minutes and started without the sample plate to preheat the block for a minimum of 2 minutes. During the preheating, one of the PCR plates was removed from 4 °C and placed at room temperature for 2 minutes. The plate was placed in the thermal cycler with the heated lid closed for 3 minutes. The heated plate was promptly removed and placed at 4 °C. This was repeated with the remaining plates using the additional programs. The plates were then spun at 500 *g* for two minutes to remove any condensation. The PCR tubes were removed from the PCR plate, carefully placed in 1.5 mL tubes, and spun at 21,000 *g* for 30 min at 4 °C to pellet the aggregate protein. Supernatant was carefully removed from each tube and placed in a clean, low-retention, 1.5 mL tubes. To reduce sample handling samples are not removed from this tube until isobaric labeling was completed: single-tube sample preparation (STSP) method. Illustration of the lysate to labeled mass spectrometry sample workflow can be found in **Supplementary Fig. 1**. 50 μL of Denaturation buffer (8 M Guanidine HCl, 100 mM HEPES pH 8.5, 10 mM Tris(2-carboxyethyl)phosphine hydrochloride (TCEP), 40 mM 2-Chloroacetamide, all prepared fresh or stored for single use at −80 °C and thawed immediately prior to use) was added to each sample, and the samples were placed in a heat block at 90 °C for 10 minutes to denature, reduce, and alkylate the samples. Samples were cooled to room temperature, 400 μL of cold acetone was added and the samples were placed at −20 °C overnight to precipitate protein. If a biphasic separation is observed, an addition of ddH_2_O and repeated centrifugation will be necessary to achieve normal protein precipitation.

### Single temperature thermal profiling in lysate

Workflow for iTSA and sample prep (STSP) is illustrated in **Supplementary Fig. 1**. Drug treated cell lysate was prepared by adding staurosporine at final concentration of 20 μM or H-89 ((N-[2-(p-bromocinnamylamino)ethyl]-5-isoquinoline-sulfonamide) at a final concentration of 100 nM (Tocris Bioscience, Bristol, UK) or harmine at a final concentration of 20 μM to prepared K562 lysate with a final DMSO concentration of 1%. A 1% DMSO vehicle control was prepared as well. Treated samples were incubated at room temperature for 10 minutes prior to the thermal shift. The PCR plate was treated as described above with the exception that the thermal cycler was held at a constant temperature with temperature indicated in the main text.

### Single temperature drug profiling in cells

120 million K562 cells grown as described above were suspended in 240 mL of warm RMPI media and split in two. Drug treated cells were prepared by adding staurosporine at final concentration of 1 μM, a DMSO vehicle control was prepared as well. The cells were placed at 37 °C for 15 minutes and then pelleted at 340 *g* for two minutes. Cells were washed twice in PBS then resuspended in a final volume of 3 mL. 100 μL/well, 5-replicates/condition of cell solution (~2 million cells) was utilized. The PCR plate was sealed and heated to 51 °C for 3 minutes as described above. The PCR plate was promptly flash frozen and stored at −80 °C. Samples were lysed by the addition of 100 μL of Lysis Buffer, vortexed, then sonicated for 30 seconds 3 times with a 30-second rest in between in a 4°C Bioruptor water bath sonicator. Lysate was centrifuged at 21,000 g, 4 °C, for 15 minutes and the supernatant collected followed by the STSP (**Supplementary Fig. 1**) with 200 μL of Denaturation buffer added to each sample.

### Sample preparation and TMT labeling

As illustrated in **Supplementary Fig. 1**, STSP, precipitated protein was collected by centrifugation at 21,000 *g*, −10 °C for 30 minutes. The acetone was carefully removed, and the pellets were washed two additional times by adding 150 μL 80% cold acetone and suspending with Bioruptor set at 4 °C for three 30 seconds sonication cycles on, 30 seconds off prior to centrifugation. (Note that if −10 °C centrifugation is not possible, we recommend incubation of acetone washes at −20 °C for a minimum of 30 minutes prior to 4 °C centrifugation during the acetone wash steps). Acetone washed pellets were dried prior to suspension in 100 μL of TMT-label compatible Digestion Master Mix (10% 2,2,2-Trifluoroethanol, 50 mM HEPES pH 8.5, 2 μg Lys-C (Wako product number 129-02541) and 2 μg trypsin (Sigma product number T6567)). Suspension in Digestion Master Mix was aided by using the Bioruptor set at 4 °C for three 30 seconds sonication cycles on, 30 seconds off, repeating Bioruptor program as needed until a visibly homogenous suspension was formed. Samples were digested for 16 hours at 37 °C with 1,200 rpm agitation in a Thermal Mixer C (Eppendorf, Hamburg, Germany).

Samples were isobaric labeled (Thermo Fisher Scientific PN 90119) according to the manufacturer recommended procedure. Briefly, 10 μL solution of 20 μg/μL TMT label in anhydrous acetonitrile (ACN) was added to the sample. Samples were incubated in label at room temperature for 1 hour. Labeling was quenched by addition of 8 μL of 5% (w/v) hydroxylamine and incubation for 15 minutes at room temperature. At this point, the 10 samples were combined into a single tube and 10 μL of 10% trifluoroacetic acid (TFA) was added. Samples were dried by vacuum centrifuge to remove ACN added with TMT label and then cleaned up using 3 washes on an Oasis HLB column (Waters PN 186000383). Mobile phases used were (A) 0.1% TFA aqueous and (B) 70% acetonitrile (ACN). Briefly, HLB columns were activated 1X in 1 mL B, equilibrated 2X in 1 mL A, and the acidified sample added. Sample was then washed 3X with 1 mL A and then 200 μL of B was used to elute peptides. Samples were dried by vacuum centrifuge and stored dry at −20 °C until ready for reversed-phase fractionation.

### RP/RP-MS/MS analysis

The TMT labeled tryptic peptides were fractionated on a high pH reversed-phase C18 column using an Agilent 1100 HPLC utilizing established methods^22^. Briefly, mobile phases used were (A) 10 mM ammonium formate, pH 10 water and (B) 10 mM ammonium formate, pH 10 in 80% (v/v) acetonitrile. Samples were loaded on a C18 column (Xbridge Peptide BEH C18, 130 Å, 2.5 μm, 2.1 x 150 mm, Waters, Milford, MA) at a flow rate of 300 μL/minute equilibrated in 5% B. Peptides were eluted with a gradient from 5% B to 100% over 120 minutes. Fractions were collected for 1 minute each, concatenating throughout the gradient for 24 mixed fractions of nearly equal complexity. Fractions were dried by vacuum centrifugation and stored at −80 °C.

All fractions for mass spectrometry were suspended in 20 μL of 5% (v/v) acetonitrile, 0.1% (v/v) trifluoroacetic acid and analyzed by direct injection on a Waters M-class Acquity column (BEH C18 column, 130 Å, 1.7 μm, 0.075 mm x 250 mm) at 0.3 μL/minute using a nLC1000 (Thermo Scientific). Mobile phases used were (A) 0.1% formic acid aqueous and (B) 0.1% formic acid acetonitrile (ACN). Peptides were gradient eluted from 3% to 20% B in 100 minutes, 20% to 32% B in 20 minutes, and 32% to 85% B in 1 minute. Mass spectrometry analysis was performed using an Orbitrap Fusion (Thermo Scientific). MS2 were collected at 50,000 FWHM resolution 2 x 10^5^ automatic gain control (AGC), 120 ms max fill time, with 20 s dynamic exclusion +/- 10 ppm using the data dependent mode “Top Speed” for 3 s on the most intense ions.

### Data Analysis

Analysis code can be accessed online at: https://github.com/CUOldLab/iTSA

#### MS/MS Search Parameters

All Thermo MS/MS raw files were processed using MaxQuant version 1.6.3.3^23^ with MS/MS spectra searched using the Andromeda search engine^24^ against a Uniprot human database (downloaded August 30, 2017, 42,215 entries) and with the TMT lot-specific isotopic distributions added for corrected reporter ion intensities. The search was limited to peptides with a minimum length of six amino acid residues with a maximum of two missed cleavages with trypsin/P and LysC/P specificity. Variable modifications for methionine oxidation and fixed modifications for protein N-terminal acetylation and carbamidomethyl Cysteine were added, and search tolerances were set at 10 ppm for MS1 precursor ions and 20 ppm for MS/MS peaks. Protein and peptide level false discovery rate thresholds were set at 1%. At least one unique or razor peptide was required for protein identification. Unique plus razor peptides were used for quantification.

#### iTSA Data Analysis

MaxQuant’s “proteinGroups.txt” output file was utilized for bioinformatics analysis using R (version 3.5.2)^25^; R script code can be accessed online at: https://github.com/CUOldLab/iTSA. The data was analyzed with tools from the R package, limma^26^. Briefly, the data matrix was filtered to include only high confidence identifications, quantile normalized, and then analyzed with linear modeling, two-sample tests with empirical Bayes statistics, and the Benjamini-Hochberg Procedure (BH) for multiple testing correction to control the false discovery rate^27–29^. Absolute fold change and q-value cutoffs were used to categorize significant protein groups for the “Sig_FDR.FC” and “rank” columns observed in the exported data files. Perseus (version 1.6.1.3)^30^ was used for calculating other statistical measures on the quantile normalized log2 intensity data. Kinome trees were built using the kinmap (http://kinhub.org/kinmap/) beta tool (**Figs. 1g & 2[d-f]**). Trees indicate kinase group target assignment and node diameter is proportional to significance of thermal shift.

To compare the iTSA data to published TPP and kinobead data the protein IPI accession numbers in the TPP and kinobead data were matched to UniprotKB identifiers in the iTSA data, and the tables were then merged in R (**Supplementary Data File 1**). TPP target proteins were considered significant (1) or not significant (0) if the protein fulfilled all 4 requirements as designated significant in the original publication^10^. Kinobead data was considered significant if a pIC50 was reported and passed quality control filters as determined in the original publication^12^.

#### Subsampling Data Analysis

We used the staurosporine iTSA 52°C data for subsampling data analysis. Our dataset contained 5 control samples and 5 drug-treated samples. Thus we can select sub-sets from both populations and calculate the significant changes (FDR < 0.1) we would have observed if our experimental design were comparing 2, 3, or 4 replicates. There were 5-choose-4 = 5 unique ways of sub-sampling each the control and drug treated samples, which means there were 25 (5×5) 4-replicate comparisons within our data. Likewise, for the 3-replicate or 2-replicate scenarios, there were 100 (10×10) possible sub-sampled comparisons. We sub-sampled from all possible combinations within our data and followed the same analysis procedure as described above to identify which proteins were identified as significant in each sub-sample. A protein was tallied if it appeared significant in >80% of sub-sampled comparisons, corresponding to a statistical power of 0.8 with the resulting percent significance (#significant/#tests) reported in **Supplementary Data File 1**. There were few proteins that were not significant in the 4 or 5-replicate comparisons that appeared significant in >10% of the sub-samples in the 2 or 3-replicate comparisons. This indicates that Type I errors are well controlled.

#### Thermal Gradient Data Analysis

MaxQuant’s output file was used for analysis and characterization of thermal profiles. Thermal gradient data was median normalized prior to fitting with parametric sigmoidal function. This normalization is performed to remove or reduce any systematic effects introduced in the peptide handling or labeling. Briefly, the median solubility change is calculated across all proteins for each isobaric label. The set of median solubilities is itself a thermal profile which can be fit to a curve. After the fit, a scale factor is calculated which can scale the median from a particular isobaric label to match the prediction from the best-fit curve. These scale factors should be very close to 1 (1 = no scaling); experiments with greater than 10% scale factor variance should not be evaluated. The calculated scale factor is then applied to all of the measurements quantified by that label.

The parametric model used to perform the normalization as well as to fit the normalized thermal profiles is the three-parameter log-logistic function:

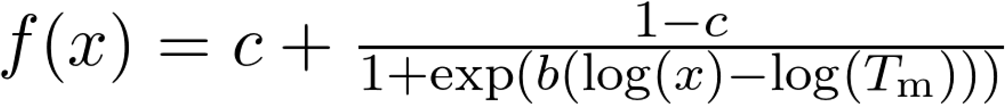

This function fits the soluble fraction relative to the lowest temperature measurement, and thus fixes the upper asymptote at 1.0, where *c* is the solubility (the lower asymptote), *T*_m_ is the melting temperature, and *b* is the slope-controlling parameter. These fits were performed in Python, using the SciPy^31^ optimization.curve_fit package (version 1.0.1) and filtered by a measure of goodness-of-fit (X^2^ ≥ 0.1). Note that this is a different model than is used in previously published studies^10^, which determine *T*_m_ with a different sigmoidal model function. To justify that this model and our sample preparation methods are comparable, we re-fit previously published data with the three-parameter log-logistic model and plotted the *T*_m_ against *T*_m_ values calculated from a new 10-temperature thermal curve using K562 cells grown in our lab (**Supplementary Fig. 4**).

#### TPP Data Analysis

The supplemental data containing staurosporine and DMSO vehicle thermal profiling data was downloaded from Savitski et al., 2014^10^ (table S4). A library of K562 *T*_m_ values was then created from this data. The *T*_m_ values for each curve fit was quality filtered by a measure of goodness-of-fit (R^2^ > 0.8) and filtered by a defined experimental range (40 < *T*_m_ < 64). The average vehicle *T*_m_ and staurosporine *T*_m_ was then calculated (n=2). The library of K562 vehicle *T*_m_ values is reported in **Supplementary Data File 1**.

### Data Availability

Raw mass spectrometry data files (Thermo raw files) and MaxQuant output files are deposited in the MassIVE repository with the primary accession code *MSV000083640*. The MassIVE dataset doi is (doi:10.25345/C55036), the URI is (http://massive.ucsd.edu/ProteoSAFe/dataset.jsp?task=0171aeef20364c1e9a22501be2d8cbdf) and the ftp location is (ftp://massive.ucsd.edu/MSV000083640).

---

**Supplementary Figure 1.**
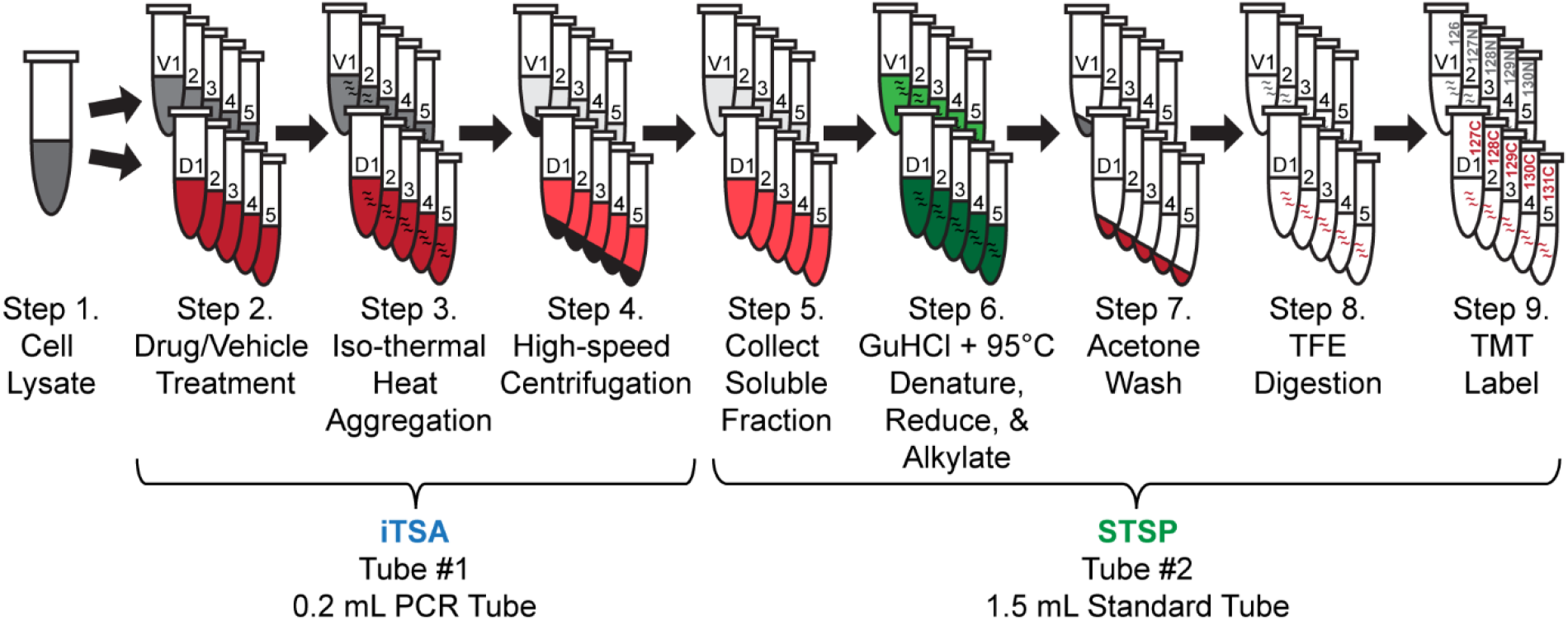
Illustrated workflow for in lysate iTSA and single-tube sample prep (STSP). Procedure focuses on minimal sample handling to reduce variance between samples. Steps are described in the figure with details for each step described in Methods.

**Supplementary Figure 2.**
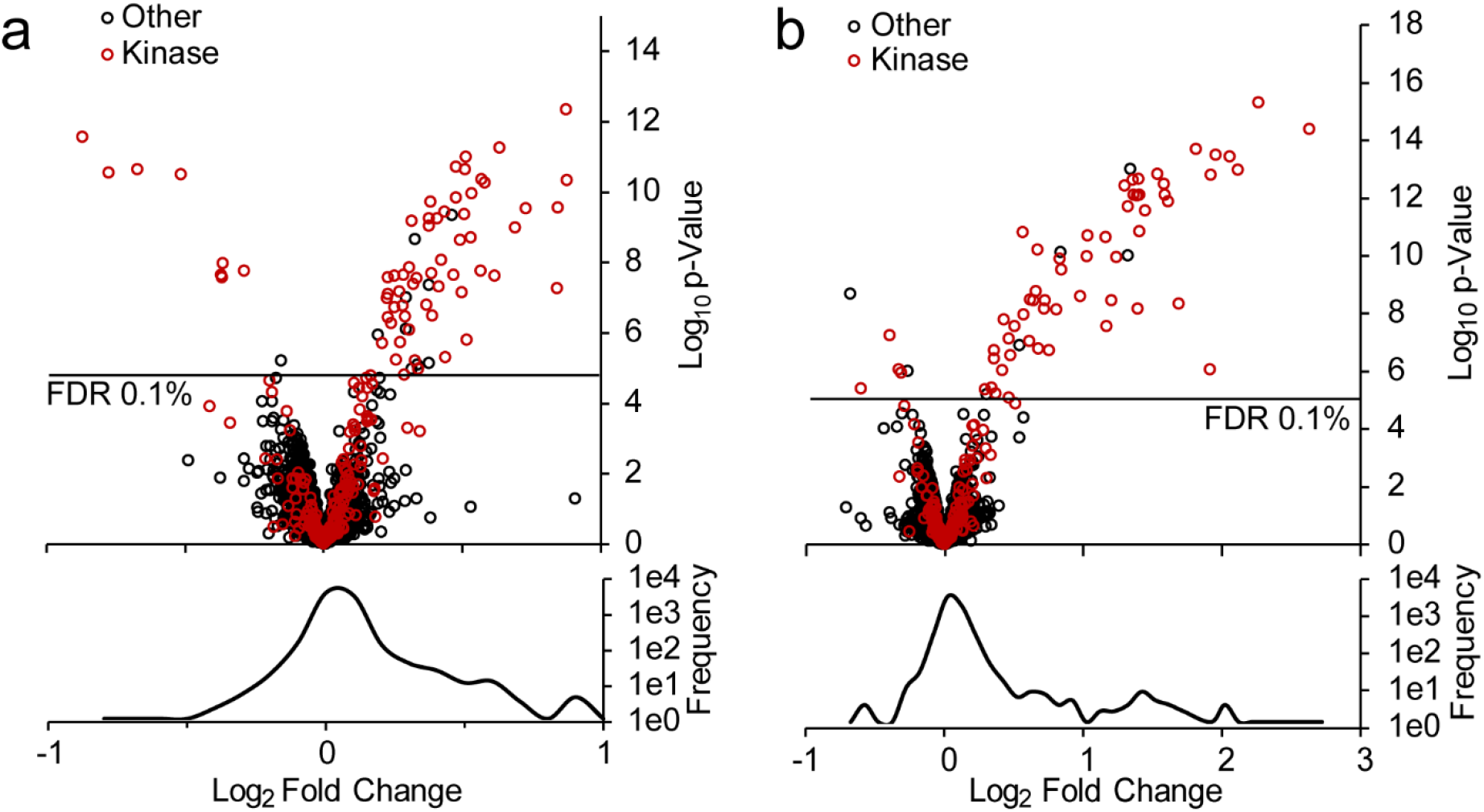
iTSA was performed at 48°C **a,** and 56°C **b,** as described in Methods using 20 μM staurosporine. Volcano plot visualization of soluble protein levels using empirical Bayes statistical analysis. Proteins identified as kinases are red. FDR = 0.1% is indicated. Frequency (aka: density/distribution) of the volcano plot x-axis are shown as well. A positive fold change represents a more stable protein in drug condition and a negative fold change represents a less stable protein in drug condition.

**Supplementary Figure 3.**
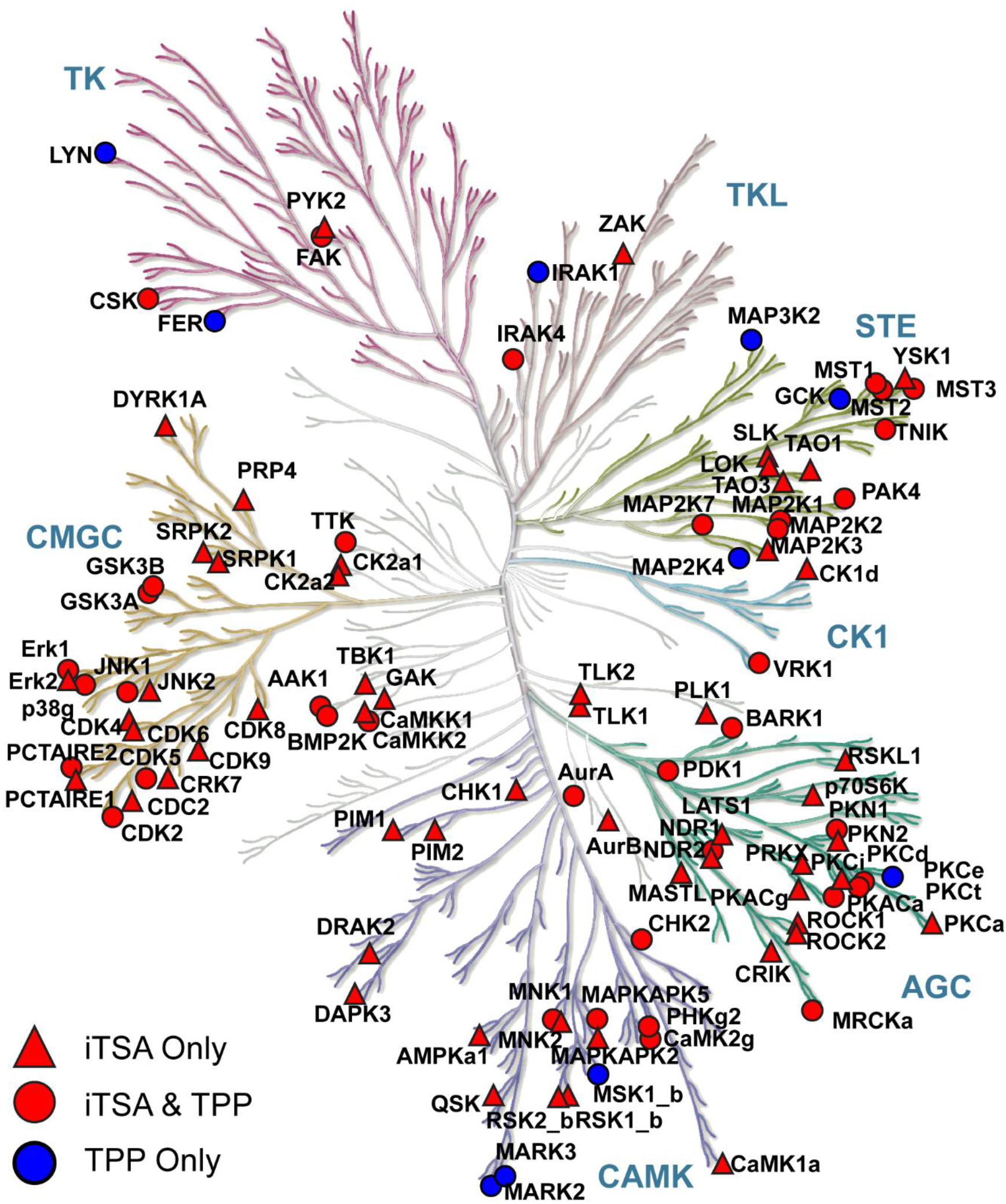
Kinome map of iTSA and TPP kinases. Blue circles represent kinases uniquely identified with TPP. Red represents kinases identified with iTSA where red triangles were uniquely identified in iTSA and red circles were targets identified by both thermal shift assays. Illustration reproduced courtesy of Cell Signaling Technology, Inc.

**Supplementary Figure 4.**
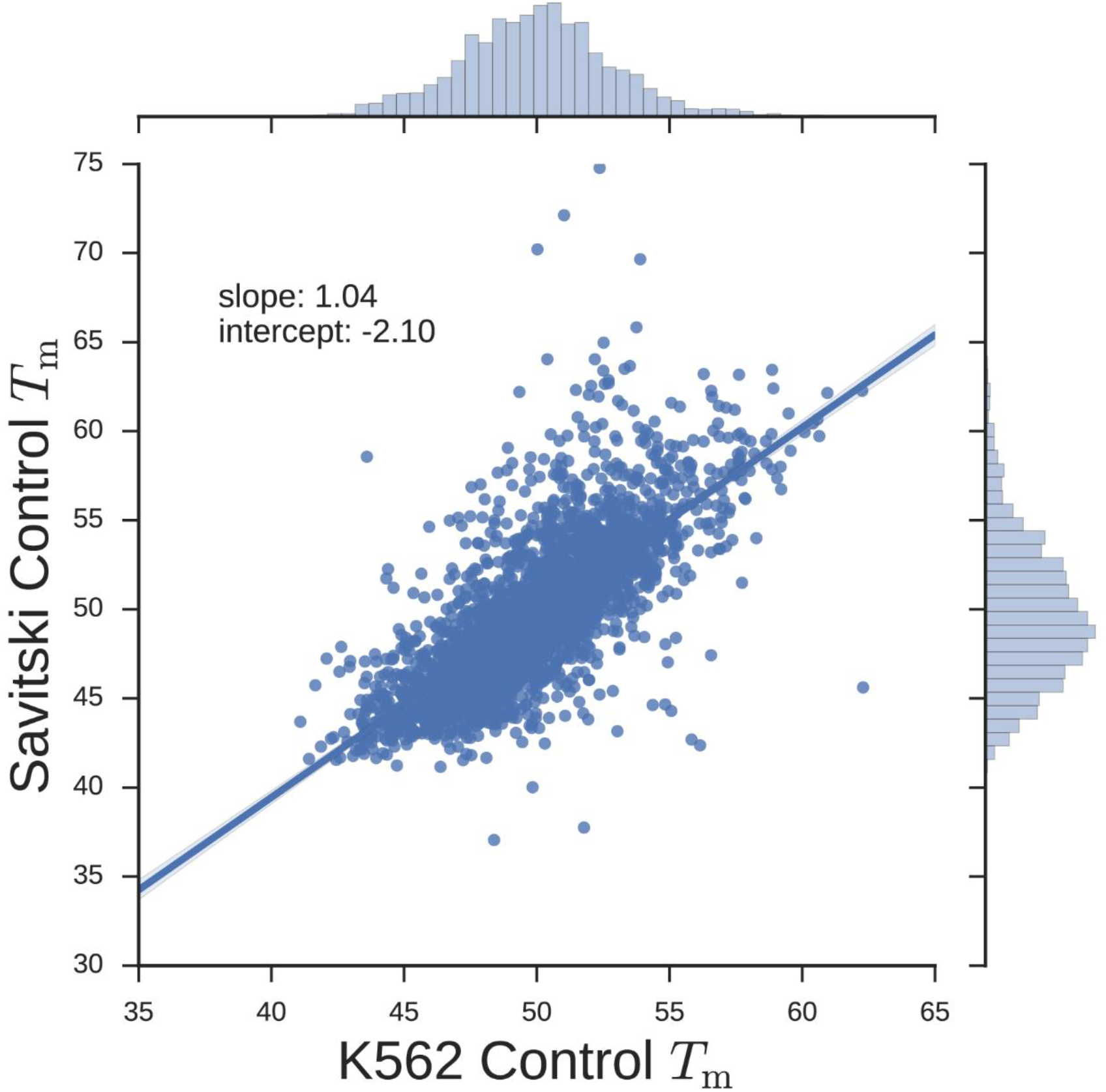
Using the three-parameter model in SciPy^31^ (Methods), we compared *T*_m_ values calculated using the previously published data^10^ to that of our full 10-temperature thermal curve performed on K562 cells. This same-software comparison illustrates that the differing sample preparation methods produce comparable curves.

